# A two-gene strategy increases the iron and zinc concentration of wheat flour and improves mineral bioaccesibility for human nutrition

**DOI:** 10.1101/2022.02.15.480518

**Authors:** Sophie A Harrington, James M Connorton, Natasha I M Nyangoma, Rose McNelly, Yvie M L Morgan, Mohamad F Aslam, Paul A Sharp, Alexander A T Johnson, Cristobal Uauy, Janneke Balk

## Abstract

Dietary deficiencies of iron and zinc cause human malnutrition globally, which can be mitigated by biofortified staple crops. Conventional breeding approaches to increase grain mineral concentrations in wheat (*Triticum aestivum* L.) have had only limited success so far due to relatively low genetic variation. Here we demonstrate that a transgenic approach combining endosperm-specific expression of the wheat vacuolar iron transporter gene *TaVIT2-D* with constitutive expression of the rice nicotianamine synthase gene *OsNAS2* has the potential to dramatically improve mineral micronutrient intake from wheat products. In two distinct bread wheat cultivars, we show that the VIT-NAS construct led to a two-fold increase in zinc to ∼50 µg g^-1^ in wholemeal flour and a two-fold increase in both zinc and iron in hand-milled white flour. In highly pure, roller-milled white flour, the concentration of iron was enhanced three-fold to ∼25 µg g^-1^. A greater than three-fold increase in the level of the natural plant metal chelator nicotianamine in the grain of VIT-NAS lines was associated with improved iron and zinc bioaccessibility in white flour. The growth of VIT-NAS plants in the greenhouse was indistinguishable from untransformed controls. We conclude that the effects of each gene cassette are additive in altering the total concentration and distribution of iron and zinc in wheat grains. This demonstrates the potential of a transgenic approach to enhance the nutritional quality of wheat well beyond what is possible by breeding approaches in order to alleviate dietary mineral deficiencies.

## INTRODUCTION

Low dietary intake of the essential mineral micronutrients iron and zinc from staple crops contributes to the global burden of malnutrition. Iron deficiency is the leading cause of anaemia worldwide, particularly affecting young children and adult females (1). In children, iron deficiency can lead to stunted development, while in adults it can severely hinder economic productivity and greatly enhances the risk of maternal death in childbirth (2). Zinc deficiency affects all age groups and genders, manifested in stunting and reduced immunity to infectious diseases (3). In some South Asian and sub-Saharan countries with cereal-dominated diets, over half of the population is predicted to be zinc-deficient (4).

Bread wheat (*Triticum aestivum* L.) is a global staple which provides between 20 and 25% of calories worldwide (5). Yet a combination of uneven nutrient distribution and associated anti-nutrient factors within the grain make wheat a suboptimal source of iron and zinc in human diets (6). Iron and zinc are predominantly located in the aleurone tissue and embryo (7-10). These parts of the grain are removed during industrial-scale roller milling of white flour, leading to the loss of 65-75% of the iron and zinc present in the whole grain (6, 11). Additionally, the bran fractions enriched in aleurone, pericarp, and embryo tissues contain high amounts of phytate (myoinositol-1,2,3,4,5,6-hexakisphosphate), an anti-nutrient which inhibits iron and zinc bioavailability (12, 13). Efforts to improve iron and zinc concentrations in wheat, therefore, must deal with the complementary issues of low levels of mineral micronutrient in the starchy endosperm, and low bioavailability in the outer cell layers comprising the bran.

Attempts have been made to leverage natural variation in wheat germplasm to increase grain iron and zinc concentrations by conventional breeding. A recent analysis of mineral micronutrients in the Watkins panel of wheat landraces identified variation of 24 to 49 µg g^-1^ zinc in the wholemeal flour, and 8 to 15 µg g^-1^ zinc in white flour (14). Similar levels of variation in zinc have been found in panels of Indian-adapted wheat varieties and CIMMYT germplasm (15-17). Variation in iron concentrations is also seen within wheat germplasm, particularly landraces and wild relatives; a panel of 170 elite, landrace, and wild relative varieties had wholemeal iron concentrations ranging from 25 to 56 µg g^-1^ iron (18). This existing genetic variation has been used to improve wholegrain micronutrient levels in elite wheat varieties. For example, introgression of the transcription factor *NAM-B1* from *T. turgidum* ssp. *dicoccoides* into bread wheat led to an increase in iron and zinc content of 18 and 12%, respectively (19, 20). However, while genetic variation in the wholegrain iron concentration exists within germplasm stocks, to our knowledge no significant natural variation has been observed for iron in white flour, nor for increased iron bioavailability (21). As a result, research effort has turned towards implementing transgenic and cisgenic approaches to improve iron and zinc levels and their bioavailability (6, 22).

Early work towards enhancing micronutrient levels was carried out in rice, exploiting genes involved in iron uptake, transport and storage. *NICOTIANAMINE SYNTHASE* (*NAS*), encoding the enzyme that synthesizes the metal chelator nicotianamine (NA), plays a key role in iron uptake and mobility of divalent metal ions in plants (23). Overexpression of either of the three individual *NAS* genes in rice not only led to increased concentrations of NA, iron, and zinc in rice grains, but also improved iron bioavailability (24-27). An alternative approach to improving grain micronutrient content focussed on the iron storage protein ferritin. Endosperm-specific expression of the soybean *FERRITIN* gene in rice led to an increase in grain iron concentration by two-to-three fold (28).

Initial efforts to biofortify wheat flour with iron and zinc built on the findings in rice, targeting similar candidate genes. Endosperm-specific expression of a *FERRITIN* gene from wheat or bean in bread wheat led to a ∼60% increase in total grain iron (29, 30), however X-Ray Fluorescence imaging showed that iron accumulated in the crease of the grain, not in the endosperm (31). High expression of the rice *OsNAS2* gene in wheat under a constitutive promoter led to 40 – 100% more iron and 60 – 250% more zinc in whole grains (30). Field trials of wheat lines generated in a different study but with a similar *OsNAS2* gene cassette showed increases of up to 30% more iron and up to 50% more zinc in whole grain and white flour. Whereas improvements in the mineral micronutrient concentrations varied from year to year and in different field sites, the lines showed a robust >200% increase in NA, as well as improved iron bioavailability (32). Another strategy to modify grain iron was demonstrated in wheat by overexpressing the wheat *VACUOLAR IRON TRANSPORTER 2* gene (*TaVIT2-D*) under the wheat endosperm-specific *HIGH MOLECULAR WEIGHT GLUTENIN-D1* promoter (33, 34). This resulted in redistribution of iron to the endosperm region adjacent to the embryo, with a consistent ∼200% increase in iron in hand-milled white flour and ≥250% increase in highly pure, roller-milled white flour fractions, from 8 to 20 µg g^-1^ (6, 33).

Here we report on the results of combining endosperm expression of the *TaVIT2-D* gene with constitutive expression of the *OsNAS2* gene in hexaploidy bread wheat. Transformation with the so called VIT-NAS construct in two genetically distinct wheat cultivars led to significant increases in both iron and zinc concentrations in white and wholemeal flour, with additive effects of each gene cassette and minimal impacts on plant growth. NA levels were up to ten-fold higher in the VIT-NAS lines compared to control, correlating with increased iron and zinc bioaccessibility.

## RESULTS

### The VIT-NAS construct drives high levels of *TaVIT2* and *OsNAS2* expression in grain and leaf tissue

To increase the total amount of iron and zinc in the wheat grain, while maximizing the concentration in the endosperm, we generated a construct, referred to as VIT-NAS, combining the *HMWG::TaVIT2-D* cassette (33) with the *ZmUBI1::OsNAS2* cassette (35) in a binary T-DNA vector with the hygromycin resistance marker (Figure 1A). Transformation of the VIT-NAS construct was independently carried out in both the cultivar Fielder, nowadays only used for research, and the cultivar Gladius, a drought-resistant variety grown in Australia. Following selection for hygromycin resistance, a total of thirteen independent transformants were obtained in cv. Fielder, with copy numbers ranging from one (in two transformants) to more than twelve. Iron staining of grains was used to select five independent transformants alongside a null transformant for further characterisation in homozygous lines of the T_3_ generation (Table 1). In cv. Gladius, 37 independent transformation events were obtained of which three independent lines with a single insert of the T-DNA— BD1-T, BD4-T, and BD7-T, alongside their null-segregant siblings (NS)— were selected for analysis in the T_1_ generation.

**Table 1:** VIT-NAS independent transformants in cv. Fielder and controls.

| Line | Copy Number (2n) |
| --- | --- |
| A1 | 0 |
| A10 | 2 |
| B4 | 4 |
| B12 | ~12<sup>a</sup> |
| B14 | 6 |
| B71 | >20<sup>a</sup> |
| TaVIT2<sup>b</sup> | 8 |
| 22-15<sup>b</sup> | 0 |

<sup>a</sup> Quantification accuracy declines at high copy numbers.
<sup>b</sup> TaVIT2 is line 22-19 as described in (33) and 22-15 is a null transformant from the same study.

**Figure 1:**
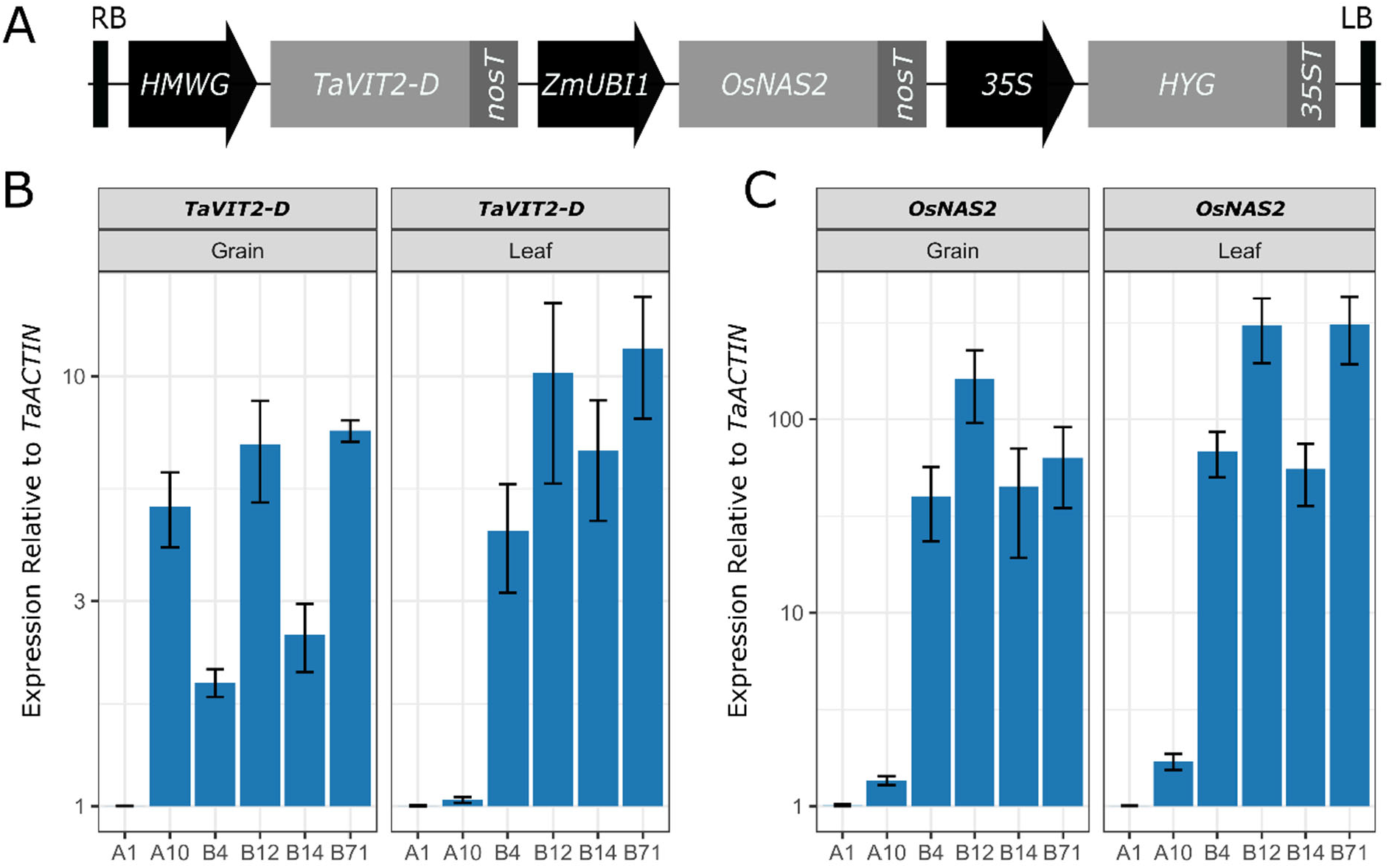
Expression of *TaVIT2-D* and *OsNAS2* in the VIT-NAS lines. A, Diagram of the transfer DNA construct (not to scale): RB, right border; *HMWG, HIGH-MOLECULAR-WEIGHT GLUTENIN-D1-1* promoter; *TaVIT2-D*, wheat (*Triticum aestivum*) *VACUOLAR IRON TRANSPORTER2-D* gene; *nosT*, (bacterial) *nopaline synthase*terminator; *ZmUBI1*, maize (*Zea mays*) *UBIQUITIN* promoter; *OsNAS2*, rice (*Oryza sativa*) *NICOTIANAMINE SYNTHASE 2* gene; *35S*, Cauliflower Mosaic Virus 35S promoter; *HYG*, hygromycin resistance gene; *35ST*, Cauliflower Mosaic Virus 35S terminator; LB, left border. B-C, RT-qPCR expression of (B) *TaVIT2-D* using primers specific for the transgene and (C) *OsNAS2* in grain (left) and flag leaf (right) tissue at 21 days post-anthesis. Expression levels calculated relative to *TaACTIN* for null transformant (A1) and transgenic (A10, B4, B12, B14, B71) lines. Error bars represent the standard error of three biological replicates for each line.

Expression analysis by RT-qPCR confirmed high levels of *TaVIT2-D* and *OsNAS2* expression in the cv. Fielder VIT-NAS lines compared to the null line A1 (Figure 1B, C). One transformant line, A10, had very low expression of *OsNAS2* in both the grain and leaf tissue compared to the other VIT-NAS lines, indicating that the gene is not properly transcribed in this line. The expression of the *TaVIT2-D* transgene in A10 is high in the grain and low in leaves, as expected from the endosperm-specific *HMWG* promoter. Unexpectedly, the other VIT-NAS lines had elevated expression levels of the introduced *TaVIT2-D* gene in leaf tissue. We speculate this may be due to insertion of several copies of the T-DNA in tandem, allowing the *ZmUBI1* or *35S* promoters to influence *TaVIT2-D* expression (see Discussion).

### Combined overexpression of *TaVIT2* and *OsNAS2* does not affect plant growth

To investigate whether expression of the VIT-NAS construct affects plant growth, we measured multiple plant growth and yield parameters for both the cv. Fielder (Fig. 2, Supp. Fig. 1) and cv. Gladius transformants (Supp. Fig. 2). In general, no significant differences were seen between the null transformant A1 and the VIT-NAS lines (Fig. 2B, Supp. Fig. 1 and 2). Some individual lines had a small but significant difference in plant height, with cv. Fielder line A10 shorter than the null segregant, while line B12 was significantly taller (p < 0.05, Dunnett Test against A1; Fig. 2). A single transformant from cv. Gladius, BD1-T, had lower tiller number than the corresponding null sibling line (p < 0.001; Supp. Fig. 2). The average grain area and length was significantly increased in B14 (p < 0.01 and 0.001, respectively; Dunnett Test against A1; Supp. Fig. 1). The remaining lines and traits, including harvest index, grain yield per plant, and thousand grain weight, showed no significant difference against the controls (Fig. 2B, Supp. Fig. 1 and 2).

**Figure 2:**
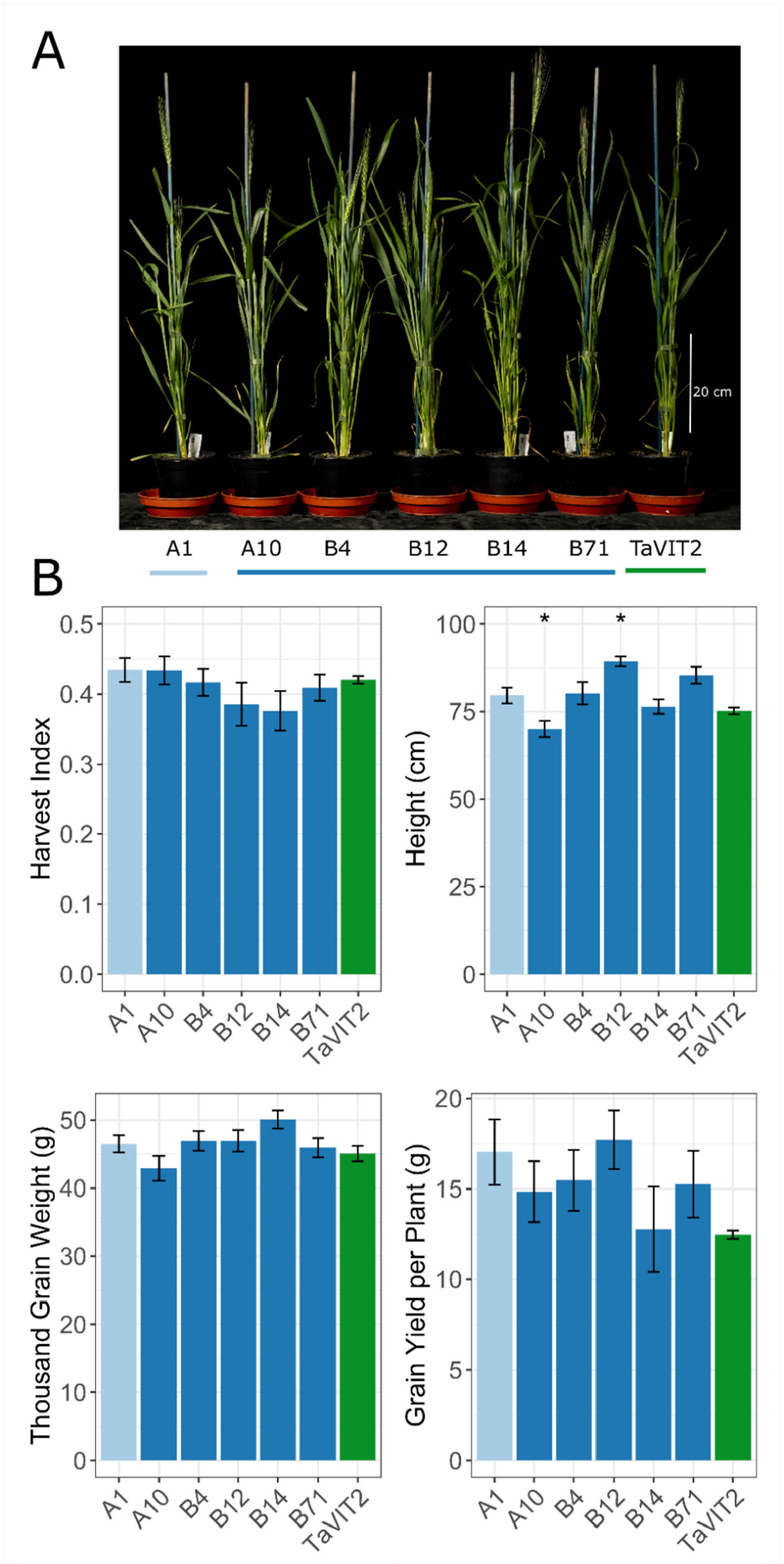
The VIT-NAS construct does not affect plant growth. A, Representative individual wheat plants of the VIT-NAS T_3_ generation at five days after anthesis. B, Plant growth parameters including harvest index, height, thousand grain weight, and grain yield per plant in the null transformant (A1; light blue), VIT-NAS (A10, B4, B12, B14, B71; dark blue), and TaVIT2 (green) lines (* p < 0.05, Dunnett Test against A1). Error bars are the standard error of five biological replicates.

### The VIT-NAS lines have increased concentrations of iron and zinc in white and wholemeal flours

To quantify the levels of iron and zinc in the VIT-NAS grains, grains were hand-milled to obtain wholemeal flour, which was sieved to obtain a crude white flour fraction. Concentrations of key micronutrients were measured using Inductively Coupled Plasma-Optical Emission Spectroscopy (ICP-OES). All VIT-NAS lines, in both cv. Fielder and cv. Gladius had approximately 2-fold higher concentrations of iron in the white flour compared with the controls, which was statistically significant in all Gladius lines and in three of the five Fielder lines (p < 0.05, Student’s t-test; Fig. 3A and Supp. Fig. 3). The two-fold increase in iron in white flour is similar to that seen in lines transformed with *HMWG::TaVIT2* only (Fig. 3A and Ref. 23). In contrast to the TaVIT2 line, all cv. Fielder VIT-NAS lines had significantly higher concentrations of zinc in the white flour fraction compared to the control lines (p < 0.05, Student’s t-test; Fig. 3A). Similarly, two of the three cv. Gladius VIT-NAS lines had significantly higher white flour zinc concentrations (p < 0.05, Student’s t-test; Supp. Fig. S3). Therefore, the increase in zinc can be attributed to overexpression of the *OsNAS2* gene in the whole plant.

**Figure 3:**
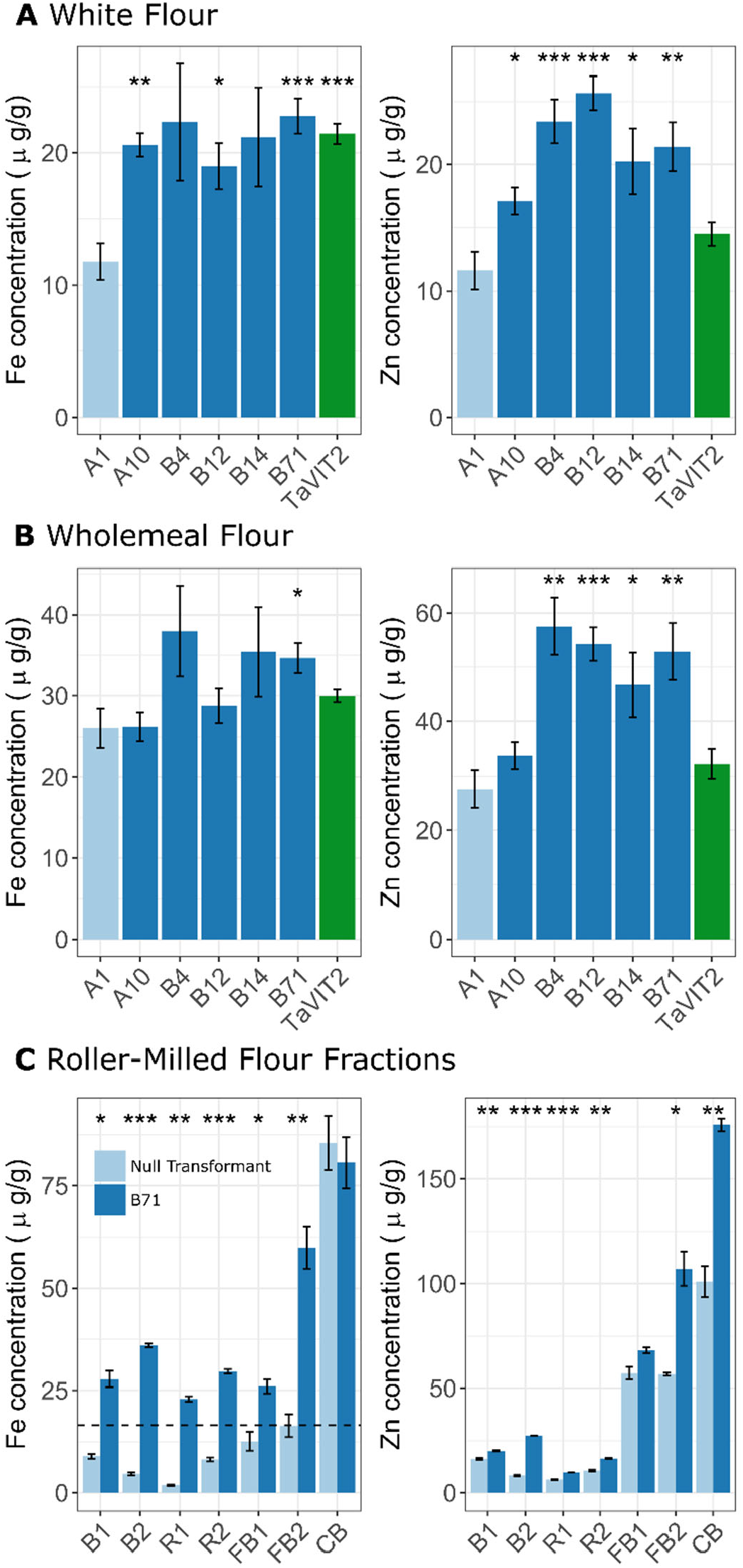
The VIT-NAS lines have increased iron and zinc concentrations. A and B, Iron and zinc concentrations measured by ICP-OES in white flour (A) and wholemeal flour (B) of the null transformant (A1, light blue), VIT-NAS (A10, B4, B12, B14, B71; dark blue), and TaVIT2 (green) lines in the homozygous T_3_ generation. Error bars represent the standard error of five biological replicates. Student’s t-test against the null (A1); *, p < 0.05, **, p < 0.01, ***, p < 0.001. C, Iron and zinc concentrations in roller-milled fractions of grain from a null transformant (light blue) and VIT-NAS (B71, dark blue). B1, first break; B2, second break; R1, first reduction; R2, second reduction; FB1, first fine bran; FB2, second fine bran; CB, coarse bran. Dashed line represents the minimum requirement for iron fortification in white flour in the UK (16.5 µg g^-1^). Error bars represent the standard error of three technical replicates. Student’s t-test between null and B71 for each fraction; *, p< 0.05, **, p < 0.01, *** p < 0.001.

In wholemeal flour, only one VIT-NAS line, B71, had significantly more iron than the control (p < 0.05, Student’s t-test; Fig. 3B). Wholemeal zinc concentrations were increased approximately two-fold in all VIT-NAS lines except A10 (p < 0.05, Student’s t-test; Fig. 3B). As noted before, the expression level of *OsNAS2* in the A10 line is low (Fig. 1B) and thus the A10 line is equivalent to the TaVIT2 line. Correspondingly, wholemeal zinc levels in both A10 and TaVIT2 are not significantly higher than in the control line. This further demonstrates the specific effect of *OsNAS2* overexpression on increasing the grain zinc concentration.

To obtain cleaner white flour fractions, grains from several plants of the VIT-NAS line B71 and a null transformant were milled using a laboratory-scale roller mill. Iron and zinc concentrations are normally between 5 – 10 µg g^-1^ in Break (B) and Reduction (R) white-flour fractions in industrially milled grain (6), similar to the concentrations we measured in the corresponding white-flour fractions of the null transformant (Fig. 3C). In the VIT-NAS lines, the iron concentration was significantly increased by three to seven-fold, to 23 – 35 µg g^-1^, and was also significantly increased in the two Fine Bran (FB) fractions (p < 0.05, Student’s t-test; Fig. 3C). A less dramatic but significant increase in the zinc concentration was found in the white-flour fractions as well as a two-fold increase in zinc in FB2 and coarse bran (CB). The increased zinc levels in VIT-NAS flour contrast with previous analysis of the TaVIT2 line, which did not have significantly increased zinc in the roller-milled white-flour fractions (12). This emphasizes both the substrate specificity of the iron TaVIT2 transporter (23) and the key role of *OsNAS2* expression in increasing zinc in wheat flours.

### The distribution of iron and zinc within the grain is altered by both the *TaVIT2* and *OsNAS2* transgenes

Analysis of hand-milled and roller-milled flour indicated that the VIT-NAS grains accumulate more iron in the tissue which comprises the white flour (endosperm). To further investigate the spatial distribution of iron within the grain, we carried out Perls’ staining on cross sections of mature grains. Similar to the TaVIT2 line (34), grains from VIT-NAS lines accumulated iron in the central region of the endosperm, between the maternal vascular bundle and the embryo (Fig. 4A, B). Iron accumulation in the endosperm of TaVIT2 lines is correlated with a decrease in the iron concentration in the aleurone (34), which can be seen by Perls’ staining in whole grain cross sections as a lack of a blue outline (compare A1 and TaVIT2 in Fig. 4A). By contrast, in the VIT-NAS grains from line B4, B12, B14 and B71, iron is retained in the aleurone tissue, although variation of staining intensity suggests the iron concentration may be less than in control grains.

**Figure 4:**
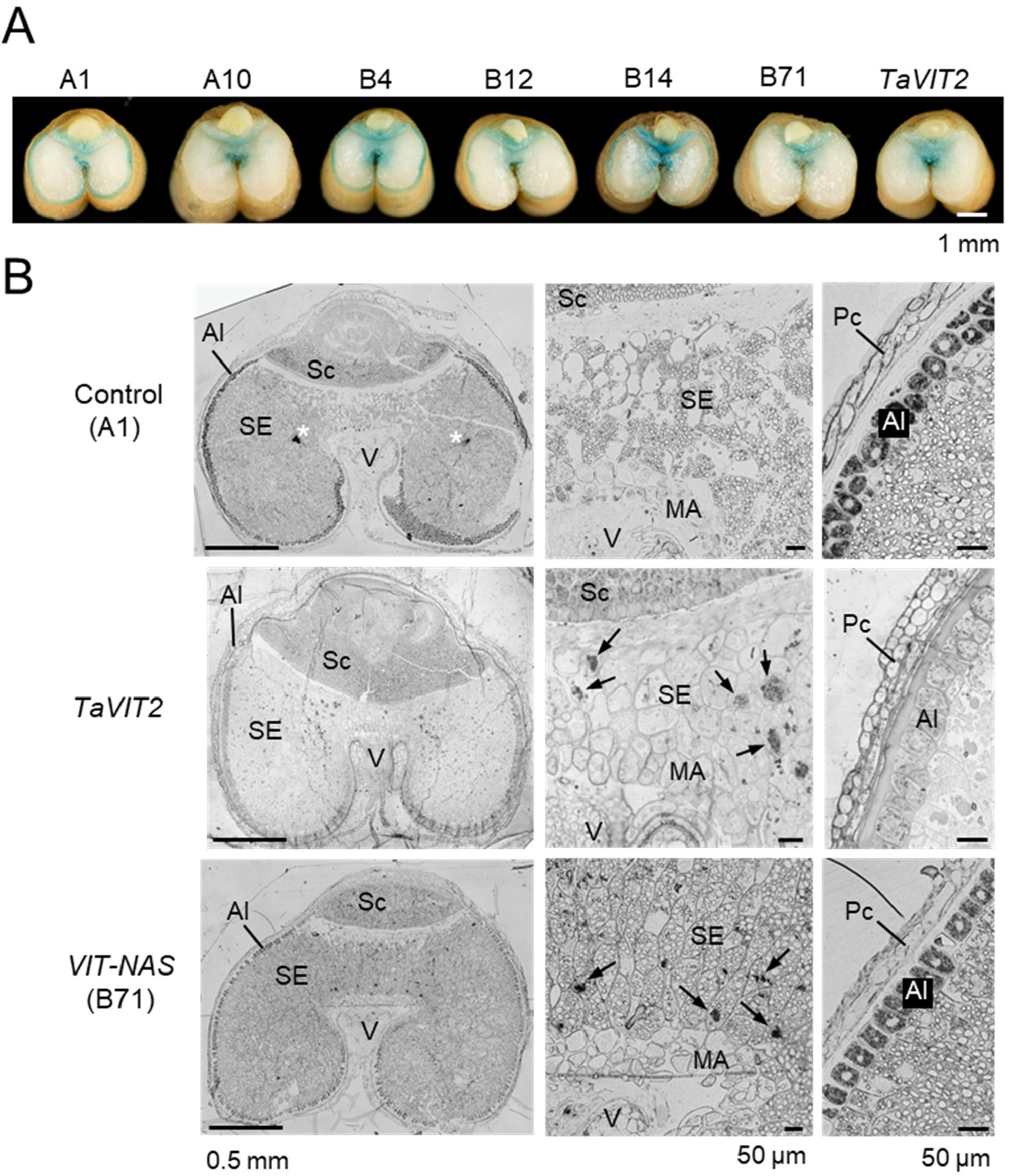
*TaVIT2* and *OsNAS2* affect the distribution of iron in the grain. A, Cross sections of mature grains stained for iron (blue) using the Perls’ method. B, Thin sections (1 µm) of immature grains 21 days after anthesis stained for iron (black in monochrome images) using the Perls’-diaminobenzidine method. Left, cross section through the grain; middle, detail at higher magnification of the starchy endosperm between the vascular bundle and embryo; right, detail including the aleurone tissue. The images are representative of two grains taken from two different plants from the indicated wheat lines. The TaVIT2 line was previously described (34). Al, aleurone; MA, modified aleurone; Pc, pericarp; Sc, scutellum of the embryo; SE, starchy endosperm; V, vascular bundle (maternal tissue). White asterisk indicates a non-specific dye precipitate; black arrows point at iron accumulation in the vacuoles of starchy endosperm cells. Scale bars as indicated.

We also carried out Perls’ staining enhanced with diaminobenzidine on thin sections of immature grains from line B71, alongside the TaVIT2 line and the null control. Iron accumulation in globoid clusters was visible in central starchy endosperm cells in both the VIT-NAS and TaVIT2 lines, but not in the control line. Moreover, iron was visibly retained in the aleurone tissue in the B71 line, compared to a strong depletion in the TaVIT2 line (Fig. 4B).

### The VIT-NAS lines have increased iron and zinc bioaccessibility

To measure the extent to which overexpression of *OsNAS2* increases the concentration of nicotianamine, we carried out HPLC-MS on wholemeal flour samples (36). We saw significant increases in nicotianamine between three and ten-fold above the control (p < 0.01, Mann-Whitney test; Fig. 5A). The wheat line without significantly increased nicotianamine levels, A10, was also the line which had failed to induce *OsNAS2* expression (Fig. 1B). The fold increase in NA is similar to that found in grain of lines overexpressing *OsNAS2* alone, which were single copy insertion lines (32, 35). Lines with two or more T-DNA inserts, and thus more copies of *OsNAS2*, displayed only a weak trend for higher grain NA concentrations. The concentration of deoxymugineic acid (DMA), a downstream metabolite of NA also associated with mineral bioavailability, was not measured, however DMA is expected to be approximately 70% of the NA levels, as shown in extensive analysis in the OsNAS2 wheat lines (32).

**Figure 5:**
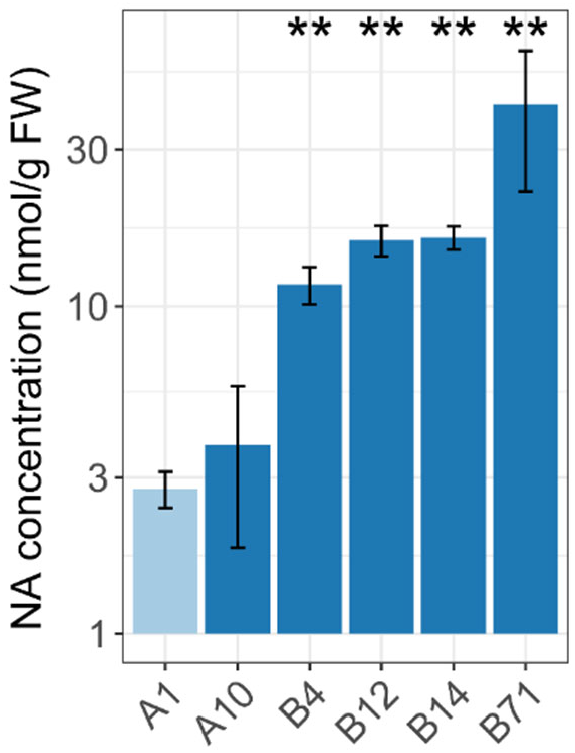
The VIT-NAS lines contain higher levels of nicotianamine. Nicotianamine concentration in grain of the control (A1, light blue) and VIT-NAS (A10, B4, B12, B14, B71, dark blue) lines. Error bars represent the standard error of three biological replicates. **, p < 0.01; Mann-Whitney test compared to the control line.

We also measured the amount of phytate, an anti-nutrient which inhibits iron and zinc bioavailability, in white and wholemeal flour. We observed no significant differences in phytate concentrations in the VIT-NAS lines compared to the control. This resulted in a two-fold increase in the iron-to-phytate molar ratio in white flour for three of the five lines tested, similar to the TaVIT2 line (Supp. Fig. 4; p < 0.05, Student’s t-test).

We then investigated the bioaccessibility of the iron and zinc levels in white flour from the VIT-NAS lines. Bioaccessibility is the quantity of soluble mineral released during simulated in vitro or in vivo digestion, relative to the total mineral in the food (37). By application of INFOGEST – a consensus, widely-accepted simulated digestion protocol (38) – we found that white flour fractions of the VIT-NAS line B71 released significantly more iron and zinc during the gastric and duodenal digestion phases relative to control flour from a null transformant line (Table 1A). In the duodenal digestion phase, iron and zinc from control white flour was largely precipitated, but more than double the amount of each mineral remains soluble in the digestate of VIT-NAS white flour (Table 1A). A similar improvement of mineral bioaccessibility was observed using a simulated gastro-intestinal digestion procedure developed for iron uptake studies in Caco-2 cells (39). Roller-milled white flour from the B71 line released 3-fold more iron and 1.3-fold more zinc compared to flour from a null transformant (Table 1B). Taken together, we see that the dramatically increased levels of nicotianamine in the VIT-NAS lines correlate with enhanced iron and zinc bioaccessibility.

**Table 1:** Iron and zinc concentrations in gastro-intestinal digests.

| | Sample | Total (μg) | Gastric digest end phase | | | Duodenum digest end phase | | |
| --- | --- | --- | --- | --- | --- | --- | --- | --- |
|  |  |  | Soluble, reagent-derived | Supernatant | Pellet | Soluble, reagent-derived | Supernatant | Pellet |
| **Fe** | Control (A1) | 1.64 | 0.03 | $0.43 \pm 0.02$ | $1.26 \pm 0.09$ | 0.63 | $0.57 \pm 0.11$ | $2.23 \pm 0.02$ |
| | NAS-VIT (B71) | 3.01 | | $0.52 \pm 0.03$ | $2.49 \pm 0.20$ | | $1.09 \pm 0.10$ | $3.20 \pm 0.01$ |
| **Zn** | Control (A1) | 2.19 | 0.10 | $2.07 \pm 0.10$ | $0.28 \pm 0.08$ | 3.06 | $1.63 \pm 0.14$ | $4.14 \pm 0.13$ |
| | NAS-VIT (B71) | 4.13 | | $3.78 \pm 0.08$ | $0.47 \pm 0.56$ | | $4.45 \pm 0.25$ | $3.54 \pm 0.09$ |

| | | Total (μM) | Released (μM) |
| --- | --- | --- | --- |
| **Fe** | Control (22-15) | 6.30 | $6.22 \pm 0.70$ |
| | NAS-VIT (B71) | 31.53 | $20.50 \pm 0.87$ |
| **Zn** | Control (22-15) | 10.80 | $5.01 \pm 0.47$ |
| | NAS-VIT (B71) | 15.96 | $8.03 \pm 0.25$ |

## DISCUSSION

Efforts to biofortify wheat must address two concerns-not only the levels of micronutrients, but also their bioavailability in human digestion. Here we show that by combining constitutive expression of the rice *OsNAS2* gene with endosperm-specific expression of the wheat *VIT2-D* gene, we can significantly increase both the quantity of iron and zinc in wheat flours and the solubility of these two mineral micronutrients in simulated gastro-intestinal digests of raw white flour. Greater solubility is likely to enhance the bioavailability of iron and zinc, although this remains to be tested in cell culture and human studies using baking products such as white and wholemeal bread.

Expression analysis of the transgenes by RT-qPCR confirmed high levels of *OsNAS2* in both leaf tissue and grain (Fig. 1C), but unexpectedly showed that the *TaVIT2-D* transgene was expressed in leaf tissue in all but one of the VIT-NAS lines (Fig. 1B). The *HMWG* promoter upstream of *TaVIT2-D* is well characterized as an endosperm-specific promoter (40), and indeed the A10 line, with a single-copy insertion of the T-DNA, showed the desired expression pattern. We hypothesise that in multi-copy insertion lines the 35S promoter of the HYG resistance marker acts as an enhancer for *TaVIT2-D* expression as noted previously in Arabidopsis studies ((41) and references therein). Leaves normally have 3.5x higher expression of the *TaVIT2* homeologs compared to grain (33), and from our phenotypic analysis there are no detrimental effects on plant growth parameters (Fig. 2, Supp. Fig. 1).

The results show that the desired effects of each expression cassette, *HMWG::TaVIT2* and *ZmUBI1::OsNAS2*, are additive. In some of the high-purity white flour fractions, we see more than five-fold improvement in the iron concentration, while the zinc concentration increased two-to-three fold (Fig. 3C). Earlier research which combined introduction of gene cassettes for endosperm-specific expression of ferritin and *UBI1*::*OsNAS2*, a strategy that was successful in rice, did not show additive effects on the mineral micronutrient content of flour and whole grain in wheat, but increases that were similar to ferritin alone (30). Possibly, entrapment of iron in the groove as a result of overexpressed ferritin (31) could have inhibitory effects on translocation of zinc into the grain mediated by NA. Another possibility is that ferritin protein in starchy endosperm cells captures only a small amount of the iron that is in transit to the embryo and aleurone cells, because the iron would need to be transported first into the plastids, across a double membrane, prior to storage in the ferritin cavity. By contrast, TaVIT2 directly transports iron from the cytosol into vacuoles for storage.

The altered distribution of iron in grain tissues of the VIT-NAS lines, caused by the unique combination of *TaVIT2* and *OsNAS2* expression, gives additional insights into the process of iron translocation during grain development, which is still poorly understood. Like the TaVIT2 line, the VIT-NAS lines accumulated iron in vacuolar globules within cells of the starchy endosperm that are located between the vascular bundle and embryo (Fig. 4B). However, unlike the TaVIT2 line, the VIT-NAS line also retained iron within the aleurone layer, similar to that observed in the null control (Fig. 4B). Isotope labelling studies combined with NanoSIMS showed that iron is trapped into the endosperm vacuoles during the nutrient-filling stage of grain development in TaVIT2 lines (34). We speculate that the retention of iron within the endosperm prevents iron from completing its movement to the outer aleurone layer. In contrast, the overexpression of *OsNAS2* in the VIT-NAS lines seems to maintain the mobilisation of iron into the aleurone layer despite the overexpression of *TaVIT2* (Fig. 4B). Perhaps the distinct expression profiles of the constitutive *ZmUBI1* and the endosperm-specific *HMWG* promoters for *OsNAS2* and *TaVIT2-D*, respectively, drive this difference. The *HMWG* promoter sequence used here is strongly activated around 14 days post-anthesis as shown by promoter:GUS studies (40). We hypothesize that expression of *OsNAS2* in the early stages of grain development promotes movement of iron (and other micronutrients such as zinc) into the grain, during which time they are transported to the outer grain layers including the aleurone cells and the embryo. Later, once the *HMWG* promoter is induced, expression of *TaVIT2* within the endosperm tissue leads to the sequestration of iron in the vacuoles of endosperm cells, as in the TaVIT2 transgenic line. The VIT-NAS line A10 supports this hypothesis, as we see iron accumulation only in the endosperm of the grain (Fig. 4A). As this line failed to overexpress the *OsNAS2* gene (Fig. 1B), it emphasizes the importance of that gene in driving the accumulation of iron in the outer layers of the grain.

Crucially, the absolute levels of zinc of at least 45 µg g^-1^ in the VIT-NAS lines are above the biofortification target of 38 µg g^-1^ (dry weight) set for whole wheat flour by HarvestPlus (42). The target for iron is 59 µg g^-1^, but this number assumes only 5% bioavailability in wholemeal flour. Depending on the cultural preference for consuming white or wholemeal flour, and the expected improvement in iron bioavailability, the iron concentrations of 25 µg g^-1^ and 30 µg g^-1^ in white and wholemeal flour, respectively, may be sufficient to achieve 30% of the estimated average requirement (EAR) of iron in the diet.

INFOGEST is fast emerging as the model digestion protocol to assess the bioaccessibility of multiple micronutrients including iron. The Glahn protocol (39) is currently used as a standard in the iron nutrition community for assessing iron release and subsequent absorption by ferritin production in Caco-2 cell culture, which has a strong positive correlation with results from human trials (37). While the Glahn protocol is facile, INFOGEST provides a robust framework for controlling many parameters including enzyme activity. In this study, improved iron release associated with the VIT-NAS flour was reflected in both digestion protocols. Phytate concentration was not significantly different between control and VIT-NAS flours (Supp. Fig. 4). For this reason, we argue that the enhanced mineral bioaccessibility seen in VIT-NAS white flour digests is a consequence of iron or zinc bound to nicotianamine which may exert preferential cation chelation compared with phytic acid. This hypothesis will be investigated by HPLC-ICP-MS in the near future.

The ultimate aim is for biofortified wheat to be grown by farmers in different parts of the world and for it to be incorporated into human diets. We found no evidence of detrimental growth effects caused by the introduction of the VIT-NAS construct (Fig. 2, Supp. Fig. 1 and 2). This will need to be further studied in field trials, but the initial data suggests that the VIT-NAS construct does not affect key agronomic traits and may thus be acceptable to farmers. It will also be important to confirm that the increased nutrient levels and bioaccessibility identified in these glasshouse trials can be replicated in the field. Promisingly, field-trials previously carried out with *OsNAS2* wheat constitutive expression lines showed increased nutrient levels and bioavailability across multiple field sites and years (32). Importantly, we show the effect of the VIT-NAS construct in two distinct wheat cultivars, Fielder and Gladius. In particular, cv. Gladius is widely grown by farmers in Australia and is known for its useful agronomic traits such as drought tolerance. The success of the VIT-NAS construct in cv. Gladius, improving micronutrient levels with no observed impact on measured plant growth traits, indicates that it could be effectively incorporated into other elite wheat varieties. A further consideration when taking a genetic-modification approach for biofortification is the regulatory landscape governing the use of such lines. It may be beneficial to develop a version of the VIT-NAS construct which uses only wheat-derived genetic sequence within the T-DNA. Research into the most appropriate wheat *NAS* gene would be required, as would replacement of the bacterial reporter genes with alternatives.

In conclusion, the stacking of *HMWG::TaVITD-2* and *UBI1::OsNAS2* results in much higher grain iron and zinc concentrations in cultivated wheat varieties than those observed in natural variety panels of wheat. Moreover, the increased NA levels are likely to compete with phytic acid to improve bioavailability of the mineral micronutrients. Future field trials are needed to confirm the results and to bulk up material for making wheat products for nutritional studies.

## MATERIALS AND METHODS

### Vector construction and wheat transformation

The VIT-NAS construct was generated by inserting the *HMWG::TaVIT2-D* cassette (33) between the T-DNA right border (RB) and *ZmUBI1*::*OsNAS2* cassette (35) in a modified pMDC32 vector backbone (43), see Fig. 1A. The *HMWG::TaVIT2-D* sequence was inserted by In-Fusion cloning (Takara Bio) in the HindIII restriction site upstream of *ZmUBI1::OsNAS2*, using primers JC172 and JC173 (Suppl. Table 1). Thus, the *TaVIT2-D* gene (*TraesCS5B02G202100*) is under the control of the *High Molecular Weight Glutenin-D1* (*HMWG*) promoter (40), resulting in ectopic expression of *TaVIT2-D* in the grain endosperm (33). The *OsNAS2* gene (Os03g0307200) is under transcriptional control of the maize ubiquitin promoter *ZmUBI1* for constitutive expression (44). The T-DNA also contains the plant-specific hygromycin resistance marker (43). The *TaVIT2* and *OsNAS2* genes were terminated by the nopaline synthase terminator (*nosT*), while the hygromycin resistance gene was terminated by the CaMV 3’ untranslated region. See Supp. Fig. 5 for a detailed map of the vector construct.

The transformation of wheat cultivar Fielder was carried out at the John Innes Centre, following the protocol detailed in (26). A zero-copy null transformant, referred to here as A1, was used as a control. At each generation, copy number analysis was carried out by iDNA Genetics (Norwich, UK) using a Taqman probe against the hygromycin resistance gene (45). Grain from the homozygous T_3_ generation was used for the micronutrient content and nicotianamine analyses reported here. The TaVIT2 line used as a control has been described in (33, 34).

The transformation of wheat cultivar Gladius was carried out at the University of Adelaide following the protocol details in (46). Copy number analysis was carried out on the T_0_ plants by quantitative real-time PCR (qPCR) using primers specific to the VIT-NAS construct (Supplemental Table 1). Individual T_0_ plants which contained single copy inserts (denoted with a “-T” ending) were selected for further characterisation in the T_2_ generation, alongside sibling null segregant lines (denoted with the “-N” ending).

### Plant growth

For all cv. Fielder experiments, seeds were pre-germinated for 48 hours at 4°C on moist filter paper. They were then sown into P96 trays containing peat-based soil (85% fine peat, 15% horticultural grit). At 21 days, the plants were transplanted into 1-liter individual pots containing Petersfield Cereal Mix (Petersfield, Leicester, UK). The plants were arranged in a randomised block design and grown in standard glasshouse conditions with 16:8 hour light:dark cycles.

For all cv. Gladius experiments, seeds were pre-germinated for 48 hours at 4 °C, after which they were germinated in P56 plug trays containing soil supplemented by 4.5 g l^-1^ Osmocote (Scotts Australia). After two weeks, plants were transplanted into 1-liter pots containing the same soil. The plants were grown in a randomised block design, initially at a constant temperature of 24°C during the day, and 18°C at night. After transplanting to individual pots, the plants were moved to an accelerated “speed breeding” growth condition (47).

### Plant phenotyping

Plant phenotyping data from the cv. Fielder plants was obtained at harvest. Plant height was measured from the soil to the tip of the highest spike, excluding awns. The entire above-ground plant was harvested and dried at 35°C for one week to obtain the total above-ground dry weight. Harvest index was calculated as the ratio of the total plant grain yield (g) to the total above-ground dry weight (g). After threshing, grain yield per plant was calculated and the thousand grain weight and grain area, width, and length were obtained using the MARVIN seed analyser (GTA Sensorik GmbH). Plant phenotyping data from the cv. Gladius plants was obtained in the same manner, with the exception that plant material was dried at 45°C for 72h to obtain the total above-ground dry weight.

### RT-qPCR

Leaf and grain tissues were sampled from individual T_3_ VIT-NAS plants (cv. Fielder) at 21 days post-anthesis and snap frozen in liquid N_2_. The snap-frozen tissue was then ground in liquid N_2_ to a fine powder and RNAs were extracted using TRIzol® Reagent (ThermoFisher). RNA concentration and purity was checked using a Nanodrop instrument (Thermo Scientific). DNase treatment was carried out using RQ1 RNase-Free DNase (Promega) before cDNAs were synthesized using the Invitrogen M-MLV reverse transcriptase.

Primers specific to the gene sequences introduced within the VIT-NAS construct were used, while previously published primers were used for the internal control *TaACTIN* (Supp. Table 2) (19). Primer efficiencies were calculated using pooled grain cDNA from lines B4, B14, B12, and B71 (Supp. Table 3). RT-qPCR reactions were performed using the LightCycler® 480 SYBR Green I Master Mix with a LightCycler 480 instrument (Roche Applied Science, UK) under the following conditions: 5 min at 95°C; 45 cycles of 10 s at 95°C, 15 s at 60°C, 30 s at 72°C; followed by a dissociation curve from 60°C to 95°C to determine primer specificity. In all cases, three technical replicates were carried out per sample and the construct-specific expression of *TaVIT2-D* and *OsNAS2* was recorded relative to *TaACTIN*.

### Flour production

For cv. Fielder, grains were hydrated to approximately 12% moisture content and milled with an IKA Tube Mill 100 for 2 minutes at 25000 RPM. The resulting wholemeal flour was passed through a 150 µm Nylon mesh to separate a crude white flour fraction from the bran. The wholemeal flour and the white flour fraction were retained for downstream analysis.

For cv. Gladius, grains were washed in a 0.1% Tween 20 solution, rinsed in distilled water, and dried at 60°C for 48 hours before milling using an IKA Tube Mill for 30 seconds at 20000 RPM. The wholemeal flour was then passed through a 200 µm Nylon mesh to obtain the white flour fraction.

For roller milled flour fractions, grain was hydrated to approximately 15% moisture content and milled using a Chopin CD1 laboratory mill. The pooled white flour for bioaccessibility assays consisted of 9.5 g of Break 1, 11.5 g of Reduction 1, 1.5 g of Break 2 and 2 g of Reduction 2 for the control flour of the 22-15 null transformant line; 9.8 g of Break 1, 11.5 g of Reduction 1, 1.5 g of Break 2 and 2 g of Reduction 2 for the B71 line.

### Elemental analysis

For cv. Fielder samples, flour was dried overnight at 65°C. Flour samples (0.1 g) were digested in 55% (v/v) nitric acid and 6% (v/v) hydrogen peroxide at 95°C for 17 hours. The acid-digested samples were 5 x diluted with ultra-pure analytical grade water. The samples were analysed using inductively coupled plasma-optical emission spectroscopy (ICP-OES, PlasmaQuant PQ 9000 Elite, Jena Analytik, Germany). Calibration was performed using 0, 0.125, 0.25, 0.5, 1, 2, 3, 4 and 5 µg g^-1^ standards of P and Mg and 0, 0.025, 0.05, 0.1, 0.2, 0.4, 0.6, 0.8 and 1 µg g^-1^ standards of Mn, Fe and Zn. Rhodium (Rh), to a final concentration of 0.1 µg g^-1^, was used as an internal standard. Hard red spring wheat reference material (National Research Council Canada) was treated in the same manner as the experimental samples and included in every run of 50 samples.

For cv. Gladius samples, the white flour fraction was analysed using ICP-OES following standard procedures at the Trace Analysis for Chemical, Earth, and Environmental Sciences (TrACEES) platform (Parkville, University of Melbourne). 1567b wheat flour was used as the standard reference material (National Institutes of Standards and Technology, MD, USA).

### Nicotianamine quantification

The concentration of nicotianamine was measured in water-based extracts using liquid chromatography-mass spectrometry (LC-MS), following the method in (36) with minor modifications. Approximately 0.5 g of wholemeal flour was mixed with 282 µl of milliQ water and 18 µl of 1 mM *N*^ε^-nicotinoyl-L-lysine (see below) which was added as internal standard. The mixture was ground for 5 min at 1000 rpm and centrifuged at 4°C at 15000 x *g* for 15 min to recover the supernatant. The pellet was extracted another 4 times with 300 µl water and the 5 supernatants pooled, then filtered through 3 kDa filters (Amicon® Ultra) at 4°C at 15000 x *g* for 60 min. The extracts and 1 µM, 10 µM, 100 µM and 200 µM standard solutions of nicotianamine (US Biological Life Sciences) each containing 83 mM *N*^ε^-nicotinoyl-L-lysine were lyophilized overnight. The residues were dissolved in 20 µl of milliQ water, 10 µl 50 mM EDTA and 30 µl of mobile phase A (1:10 ratio of 10 mM ammonium acetate to acetonitrile, pH 7.1) before being filtered through 0.45 µm polyvinylidene fluoride (PVDF) ultrafree-MC centrifugal filters (Durapore, Merck) for 10 min at 12000 x *g*. The samples were then analysed by LC-MS (Xevo TQ-S, Waters). The separation was performed using a Waters Acquity Ultra Performance LC system and a µLC column (SeQuant® ZIC®-HILIC, 150 × 1 mm internal diameter, 5 µm, 200 Å) equipped with a guard column. The flow rate of the mobile phases was set to 0.15 ml min^-1^. The gradient program was set to 100% mobile phase A for 3 min; a linear gradient to 30% A and 70% B (8:2 ratio of 30 mM ammonium acetate to acetonitrile, pH 7.3) over 7 min; 30% A and 70% B for 7 min; a gradient to 100% A for 8 min; 100% A for 10 min. The total run time for each sample was 35 min, the injection volume was 5 µl and the auto sampler temperature was 6°C, whilst the column was at room temperature. The liquid chromatography system was coupled to a time of flight-mass spectrometer (ToF-MS) with a negative electrospray source. The spray chamber conditions were set to a spray voltage of 1.5 kV, a desolvation temperature of 500°C, flow rates were 900 h^-1^ and 150 h^-1^ for the desolvation and cone gas, respectively, and the nebuliser pressure was set to 7.0 bar. The TargetLynx V4.1 (Waters Inc) software was used for quantification. The 302 to 186 *m/z* daughter transition was used to quantify nicotianamine and the 250 to 78 *m/z* daughter transition was used to quantify *N*^ε^-nicotinoyl-L-lysine.

### Bioaccessibility assays

Simulated digestion following the INFOGEST method was performed as described in (48) with minor modifications. Hand-milled white flour (0.2 g) was mixed with 600 µl of water to obtain a homogenous bolus. For the oral phase, 640 µl of simulated salivary fluid was added to the bolus, 1.5 mM CaCl_2_(H_2_O)_2_ and 156 µl of water. Human salivary amylase was omitted as oral starch digestion was not of interest. The mixture was incubated for 2 min at 37°C. For the gastric phase, the pH was adjusted to 3.0 using 0.5 M HCl, and 1.36 ml of simulated gastric fluid (as defined in Ref 46), 0.15 mM CaCl_2_(H_2_O)_2_, and 96 µl of water was added. Pepsin (Merck P7012, pepsin from porcine gastric mucosa) was added to a final concentration of 2,000 U/ml. Lipases were omitted owing to the low lipid content of the flours. Samples were incubated for 1 h at 37°C. For the duodenal phase, 1.68 ml of simulated intestinal fluid (46) was added to the gastric mixture, adjusting the pH to 7.0. To this was added 10 mM bovine bile salts (Merck B3883), 0.6 mM CaCl_2_(H_2_O)_2_, 100 U/ml porcine trypsin (Merck T4799), 25 U/ml bovine chymotrypsin (Merck C4129) and 200 U/ml porcine pancreatic α-amylase (A3176), was added. Pancreatic lipase and colipase were omitted. The intestinal phase was incubated at 37°C for 2 h.

At the end of each digestion phase, samples were centrifuged at 1000 x *g* for 5 min. Supernatants were decanted for analysis of soluble minerals; pellets were resuspended in 1 ml of analytical grade water and also analysed to obtain total values of iron and zinc for each sample. Gastric endpoint samples (2.5 ml supernatant) were acid mineralised in 16%% (w/v) HNO_3_ and 1.7% (w/v) 0.2 ml H_2_O_2_ at 95°C for 16 h, then made up to 4 ml with ultrapure water. Duodenal endpoint samples (6.4 ml supernatant) were frozen in liquid nitrogen and lyophilized prior to elemental analysis as described for flour.

Simulated gastrointestinal digestion following the Glahn protocol was performed as described in (39). Pooled, roller-milled white flour (1 g) samples were mixed with 10 ml of saline solution (140 mM NaCl and 5 mM KCl, pH 2.0) and adjusted to pH 2.0 with 1 M HCl. To this, 0.5 ml of pepsin (Sigma-P7000; 16 mg ml^-1^) was added. Samples were incubated at 37°C on a rocking platform (150 rpm) for 90 min. The pH of the samples was then adjusted to pH 5.5 using 1 M NaHCO_3_. Bile extract (Sigma-B8631; 8.5 mg ml^-1^) and pancreatin (Sigma-P1750; 1.4 mg l^-1^) was added at 2.5 ml, and the pH was adjusted again to pH 7.0. The solution was made up to 16 ml with saline solution (140 mM NaCl and 5 mM KCl, pH 7.0), and the samples were incubated at 37°C for 90 min. Samples were centrifuged at 1000 x *g* for 5 mins to separate the soluble mineral fraction and insoluble food pellet. For mineral quantification, 250 µl of supernatant was added to 14.75 ml HNO_3_ (17.5%) containing 1 mg l^-1^ Yttrium (Merck Millipore) as an internal standard. Mineral content in the samples were analysed using ICP-OES (Thermo Fisher iCAP 6300).

### Iron staining

Whole, mature wheat grains were cut with a platinum-coated razor blade and immersed in 2% (w/v) potassium ferrocyanide and 2% (v/v) HCl for 45 min, then washed with water and photographed. Grain cross sections (1 µm) were prepared as previously described, mounted on glass slides and stained for iron using the Perls’ method enhanced with diaminobenzidine (34). In brief, the slides were immersed in 2% (w/v) potassium ferrocyanide and 2% (v/v) HCl for 45 min. After washing with distilled water, samples were incubated in methanol containing 0.01 M NaN_3_, and 0.3% (w/v) H_2_O_2_ for 60 min. The slides were washed with 0.1 M phosphate buffer (pH 7.4) and incubated for 30 minutes with 1.8% (w/v) CoCl_2_.6H_2_O and 0.025% (w/v) diaminobenzidine in phosphate buffer. The slides were washed once with distilled water and left to air dry overnight and mounted with DPX mountant (Merck). The slides were imaged using a Leica DM6000 at 10x magnification.

## Acknowledgements

We would like to thank Sadiye Hayta, Mark Smedley and Wendy Harwood (John Innes Centre, JIC) for wheat transformation; Julien P. Bonneau for growth of Gladius transformants; JIC Horticulture staff for plant growth; Alison Lovegrove (Rothamsted Research) for roller milling; Marina Franceschetti (JIC) for ICP-OES; Martin Rejzek (JIC Chemistry Platform) for synthesis of nicotinoyl-L-lysine; Ana Álvarez-Fernández (Spanish National Research Council, CSIC) for advice on nicotianamine quantification; Lionel Hill (JIC Metabolomics Platform) for assistance with LC-MS; Kirstie Halsey (Rothamsted Research) for sample preparation for microscopy; Shailender Verma for iron staining of TaVIT2 sections; Eva Wegel (JIC Bio-imaging Platform) for microscopy training and assistance.

---

**Supplemental Figure 1:**
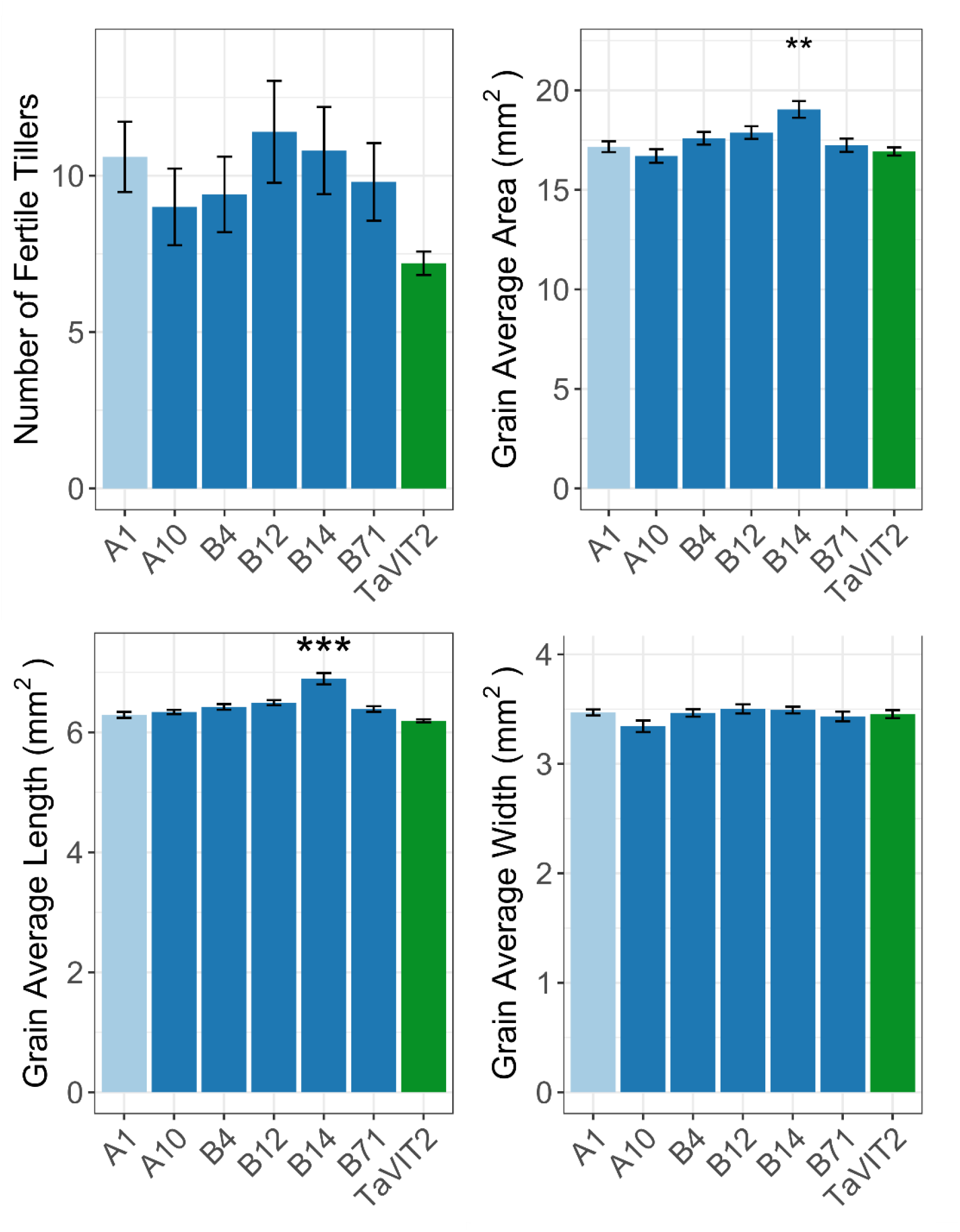
The VIT-NAS construct does not affect plant growth in cv. Fielder. Fertile tiller number and grain size parameters (area, length, and width) in the null transformant (A1; light blue), VIT-NAS (A10, B4, B12, B14, B71; dark blue), and TaVIT2 (green) lines (** p < 0.01, *** p < 0.001, Dunnett Test against A1). Error bars are the standard error of five biological replicates.

**Supplemental Figure 2:**
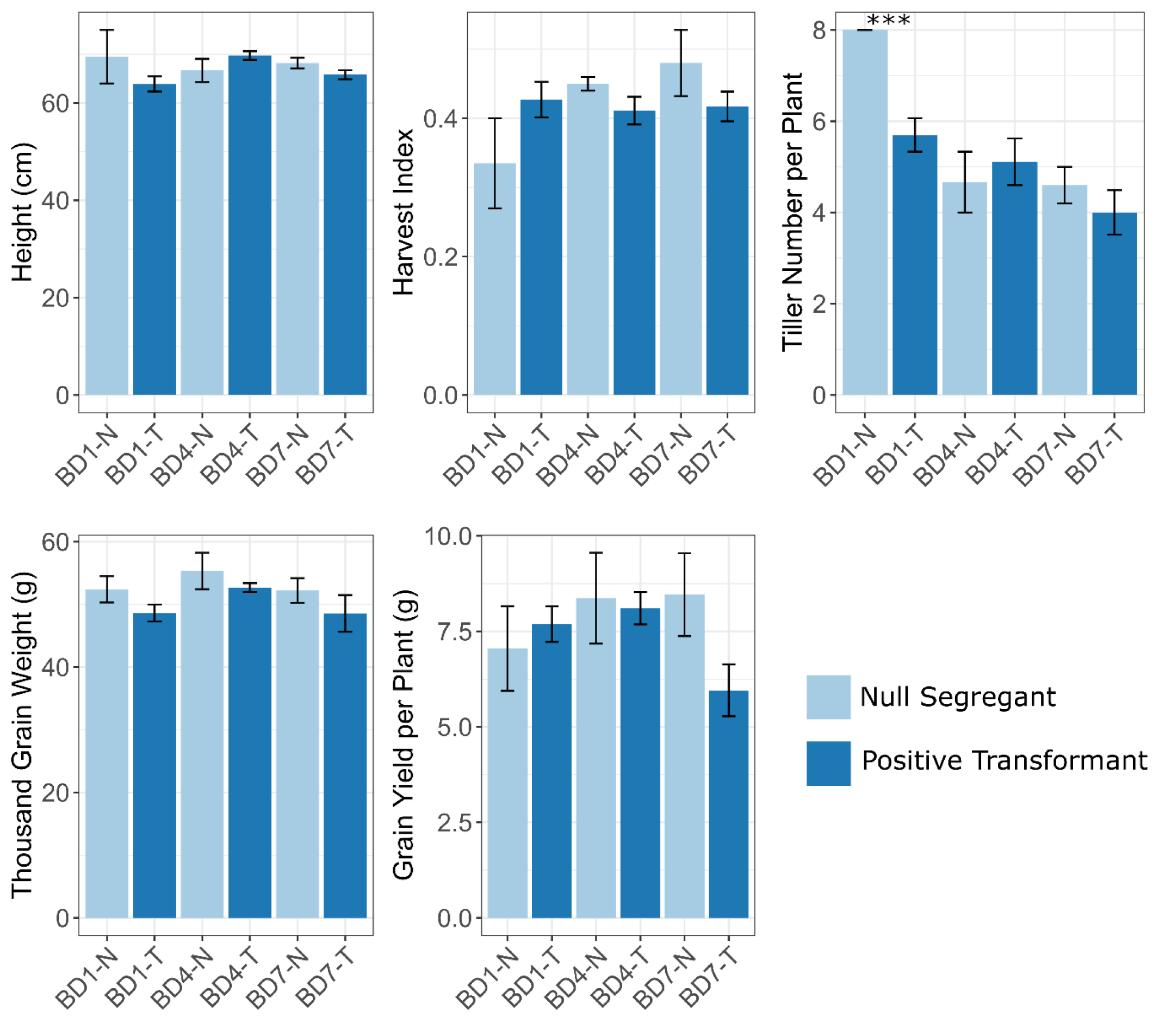
The VIT-NAS construct does not affect plant growth in cv. Gladius. Plant growth parameters including height, harvest index, tiller number, thousand grain weight, and grain yield per plant in three pairs of single-copy lines (dark blue, “-T”) and their respective null segregant sibling (light blue, “-N). *** p < 0.001, Student’s t-test. Error bars are the standard error of the biological replicates; n = 2 for BD1-N, 12 for BD1-T, 3 for BD4-N, 7 for BD4-T, 5 for BD7-N, and 7 for BD7-T.

**Supplemental Figure 3:**
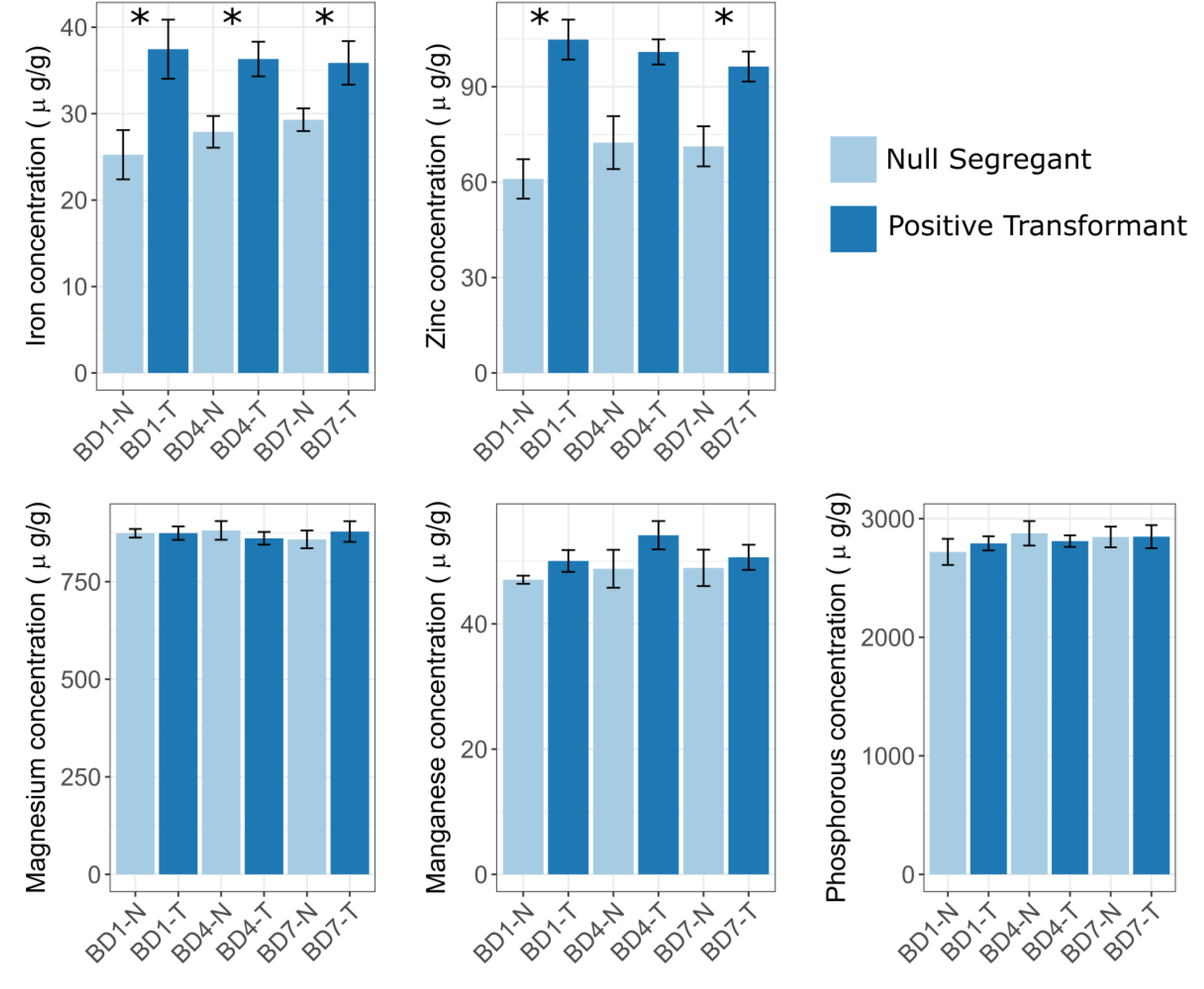
The VIT-NAS lines in cv. Gladius have increased grain iron and zinc. Micronutrient levels in white flour for the cv. Gladius VIT-NAS lines, in three pairs of single-copy lines (dark blue, “-T”) and their respective null segregant sibling (light blue, “-N). Error bars are the standard error of the biological replicates; n = 2 for BD1-N, 12 for BD1-T, 3 for BD4-N, 7 for BD4-T, 5 for BD7-N, and 7 for BD7-T. Student’s t-test was carried out for each pair against the null segregant; *, p < 0.05, **, p < 0.01, ***, p < 0.001.

**Supplemental Figure 4:**
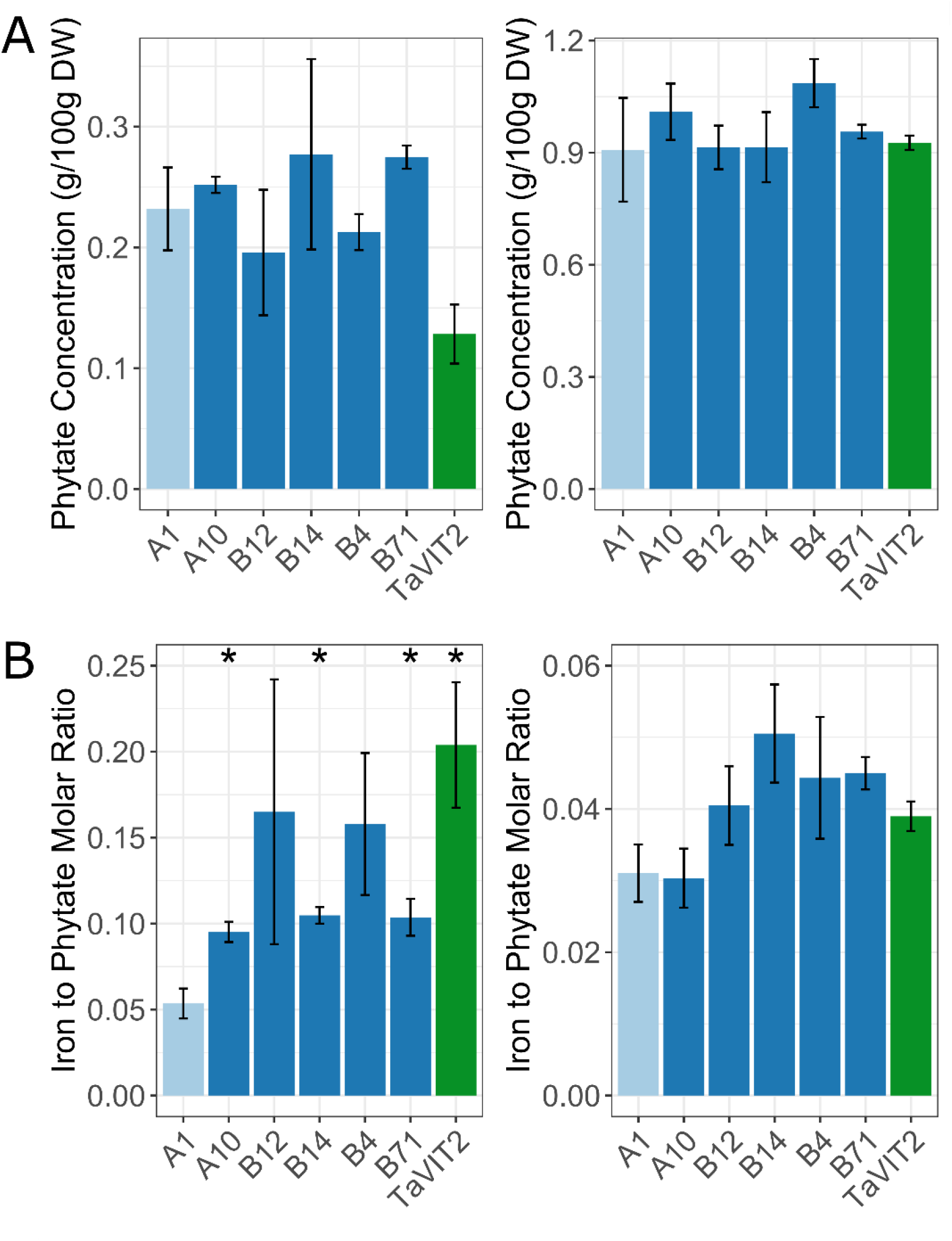
Phytate levels in the cv. Fielder VIT-NAS lines are not increased. A, Phytate levels were measured in the white (left) and wholemeal (right) flour from the null transformant (A1; light blue), VIT-NAS (A10, B4, B12, B14, B71; dark blue), and TaVIT2 (green) lines. B, The ratio of iron to phytate in the white (left) and wholemeal (right) flour for the null transformant (A1; light blue), VIT-NAS (A10, B4, B12, B14, B71; dark blue), and TaVIT2 (green) lines. Error bars are the standard error of 3 biological replicates. *, p < 0.05, **, p < 0.01, ***, p < 0.001; Student’s t-test against the null (A1).

**Supplemental Figure 5:**
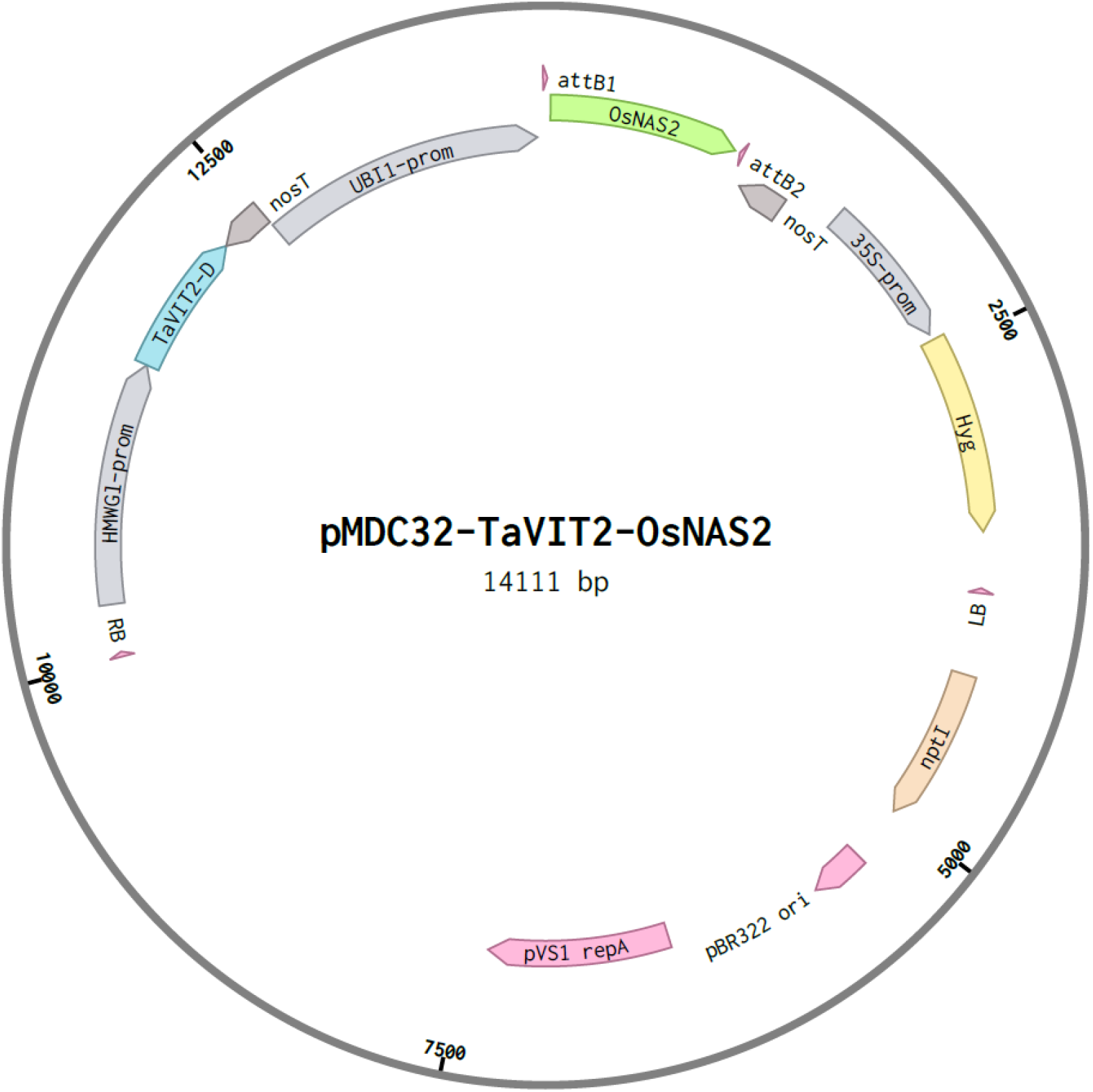
T-DNA plasmid used for wheat transformation. RB, Right Border of T-DNA; *HMWG1-prom*, promoter sequence of the High Molecular Weight *GLUTENIN-D1* gene; *TaVIT2-D*, open reading frame of the *VACUOLAR IRON TRANSPORTER 2*, D homoeologue (TraesCS5B02G202100) from wheat; *nosT*, nopaline synthase terminator; *UBI-prom*, promoter sequence of the maize *UBIQUITIN1* gene; attB1 and attB2, sequence elements (25 nt) for Gateway cloning; *OsNAS2*, open reading frame of the *NICOTIANAMINE SYNTHASE 2* gene (Os03g0307200) from rice; *35S-prom*, promoter sequence of the Cauliflower Mosaic Virus; *Hyg*, open reading frame of the plant selectable marker hygromycin phosphotransferase; LB, Left Border of T-DNA; *nptI*, open reading frame of the bacterial selectable marker neomycin phosphotransferase; pBR322 ori, origin of replication for plasmids in *Escherichia coli; pVS repA*, one of the genes for replication and stability of the plasmid in *Agrobacterium tumefaciens* (not all genes of the replicon are depicted).

**Supplemental Table 1:**
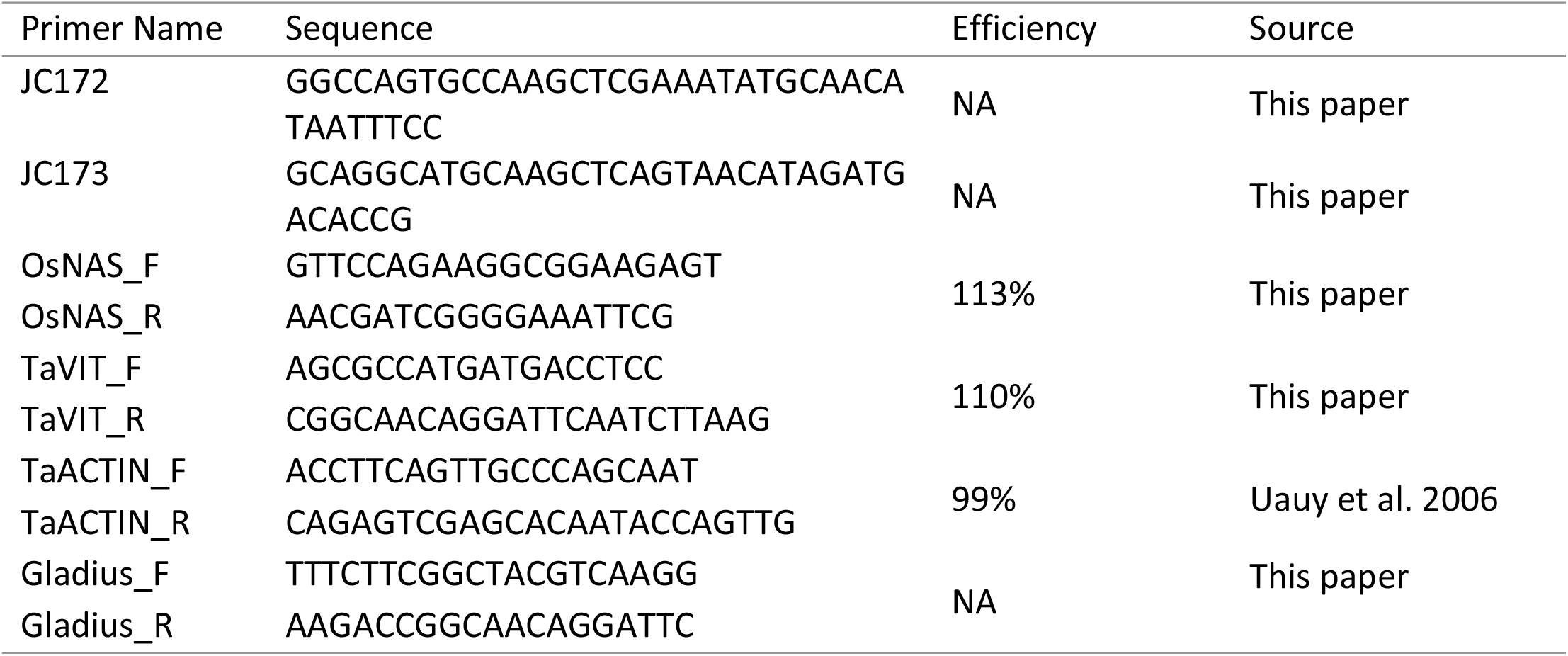
Primers used in this study. The JC172 / JC173 primer pair was used to PCR-amplify the *HMWG::TaVIT2* cassette and then insert this into the HindIII site upstream of *ZmUBI1::OsNAS2* using In-Fusion cloning, see diagram in Fig. 1A. The OsNAS, TaVIT, and TaACTIN primer pairs were used to determine the expression levels in Fig. 1B. The Gladius primer pair was used for genotyping of the VIT-NAS cv. Gladius lines.

